# Shrinking of fish under warmer temperatures decrease dispersal abilities and speciation rates

**DOI:** 10.1101/2020.10.27.357236

**Authors:** Jorge Avaria-Llautureo, Chris Venditti, Marcelo M. Rivadeneira, Oscar Inostroza-Michael, Reinaldo J. Rivera, Cristián E. Hernández, Cristian B. Canales-Aguirre

**Affiliations:** Centro de Estudios Avanzados en Zonas Áridas, CEAZA, Coquimbo, Chile; School of Biological Sciences, University of Reading, Reading, UK; Departamento de Biología Marina, Facultad de Ciencias del Mar, Universidad Católica de Norte, Coquimbo, Chile; Departamento de Biología, Universidad de La Serena, La Serena, Chile; Laboratorio de Ecología Evolutiva y Filoinformática, Departamento de Zoología, Facultad de Ciencias Naturales y Oceanográficas, Universidad de Concepción, Concepción, Chile; Centro de Investigación en Biodiversidad y Ambientes Sustentables (CIBAS), Facultad de Ciencias, Universidad Católica de la Santísima Concepción (UCSC), Concepción, Chile; Universidad Católica de Santa María, Arequipa, Perú; Centro i~mar, Universidad de Los Lagos, Camino a Chinquihue km 6, Puerto Montt, Chile; Núcleo Milenio de Salmónidos Invasores (INVASAL), Concepción, Chile

## Abstract

There is an ongoing debate as to whether fish body size will decrease with global warming and how body size changes may impact dispersal abilities and speciation rates. Although theory predicts that, when fish face warmer temperatures, they grow to smaller adult sizes, see a reduction in their ability to move, and increase their probability of speciation, evaluations of such predictions are hampered owing to the lack of empirical data spanning both wide temporal and geographical scales. Here, using phylogenetic methods, temperature, and 21,895 occurrences for 158 worldwide-distributed species of fish, we show that smaller fish have occurred in warmer waters for over 150 million years and across marine and freshwater realms. Smaller fish have historically moved the shortest distances and at low speeds. In addition, small fish display the lowest probability of giving rise to new species. Further, we found that species of fish that displayed high speeds of geographical movement and rates of size evolution experienced higher rates of temperature change in their lineage. Taking these results together, global warming predicts a future where smaller fish that have reduced ability to move over aquatic systems will be more prevalent, in turn, this will result in fewer species contributing global biodiversity.

---

A great deal of scientific research seeks the impact of human-induced global warming on Earth’s biodiversity^1–5^. Compelling evidence suggests that global warming will increase species extinction risk^6–8^, but there are hints in the literature pointing to the idea that species have several alternative strategies which might enable them to survive such adversity^2,3,9,10^. Local adaptive changes to decrease body size or tracking of suitable environmental conditions over geographic space have emerged as common responses allowing species survival, especially in fish^8–11–19^. However, it is unknown to what extent fish get smaller with warming^20^ and how these climate-induced changes in size will impact their ability to track optimal environmental conditions over aquatic systems, i.e., dispersal abilities^4,5,10^. Furthermore, the consequences that the interaction between temperature, size, and dispersal may have on speciation is less explored, even though speciation is indeed the principal buffer preventing biodiversity loss in the face of species extinction.

Based on previous knowledge, a positive association between fish size and dispersal abilities is expected given that bigger species are more efficient in consuming energy for long distance dispersals^21^. Moreover, population genetics theory postulates that organisms with high capacity to move will increase the gene flow within species and therefore predicts a low probability of population divergence and speciation^22^. Taking these predictions together, it is expected that the evolution of smaller fish under global warming (Fig. 1a) will decrease their dispersal abilities (Fig 1b) but increase the rate at which they contribute with new species to biodiversity by local genetic differentiation (Fig. 1c and d). Nevertheless, there is a big gap between theoretical expectations and evidence owing to the lack of combined data on size evolution, temperature change, species dispersal abilities and speciation rates. This patchy evidence comes by virtue of the fact that, first, the relationships between size, dispersal, and temperature change have only been evaluated across small temporal scales (i.e. decades)^12,13,17–20,23,24^, where the process of speciation cannot be observed. Second, species movement is notoriously difficult to quantify^25–27^ so that most studies use data from extremely few individuals within species, measured in recent decades^19^.

**Figure 1.**
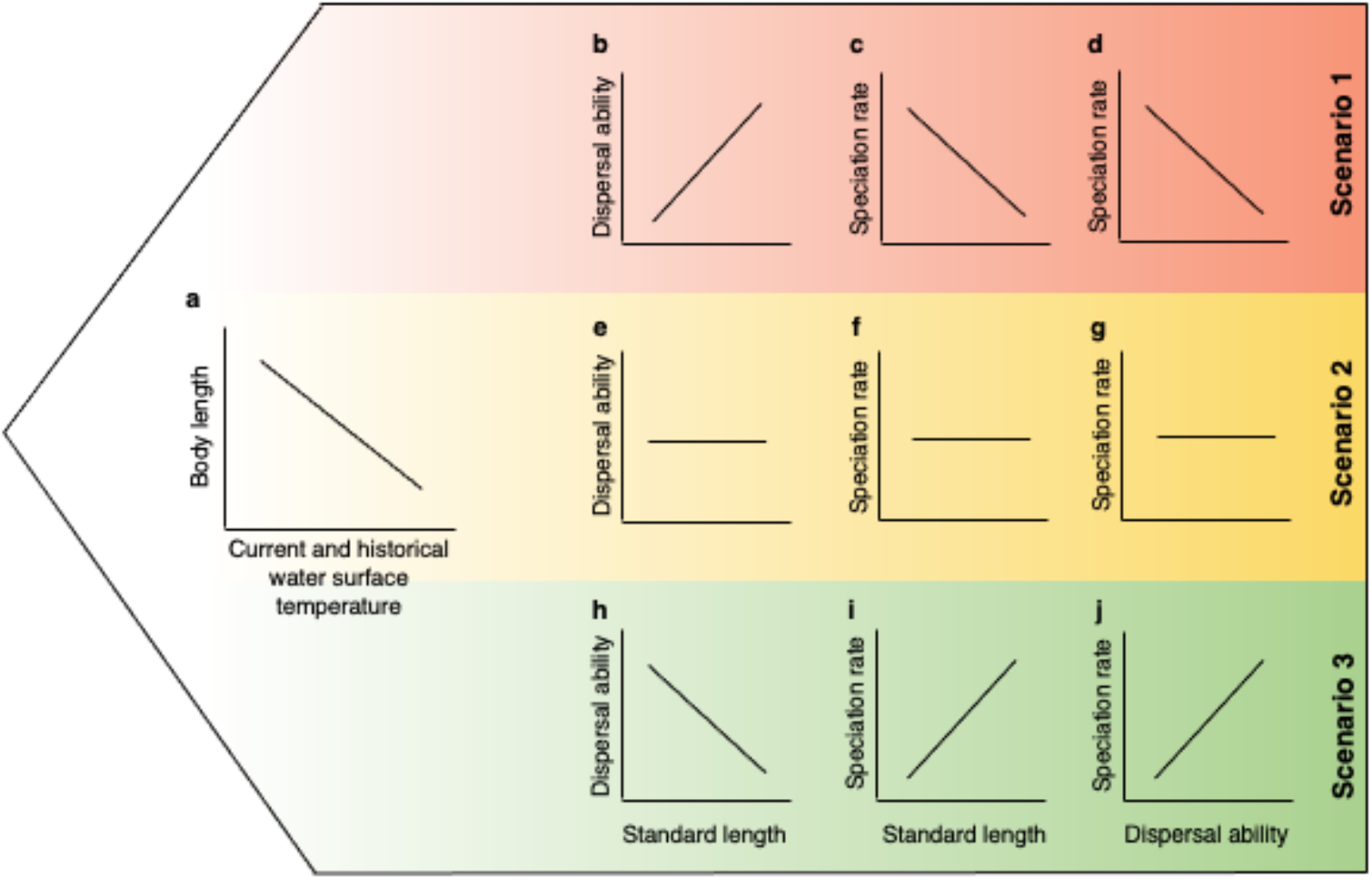
Size reductions under global warming can impact dispersal abilities and speciation rates in multiple ways. **a**, a negative relationship between standard length (SL) and water surface temperature (WST), across the phylogeny and the extant global distribution of fish, support the idea that warmer temperatures have selected small fish over million years and at wide geographical scales. **b - d,** if small fish are less likely to shift their geographic range but more prone to speciate we should observe (**b**) a positive relationship between dispersal ability and SL; (**c**, **d**)a negative effect of dispersal ability and SL on speciation rates. **e – g,**we cannot make any inference if there is no relationship between SL, dispersal abilities, and speciation rate. **h - j,** if small fish are more likely to shift their geographic range but less prone to speciate we should observe (**h**) a negative relationship between dispersal ability and SL; (**h**, **j**) a positive effect of dispersal ability and SL on speciation rates.

Here, for the first time, we test these predictions (and potential alternatives; Fig. 1) in Clupeiformes species - a highly diverse Order of fish with worldwide distribution, inhabiting the marine and freshwater realms^28^ (Supplementary Figure 1). Clupeiformes include some of the most important species for fisheries^29^, such as the anchovy *(Engraulis ringens)*, atlantic herring *(Clupea harengus)*, japanese pilchard *(Sardinops melanostictus)*, pacific herring *(Clupea pallasi)*, and the south american pilchard *(Sardinops sagax).* We evaluated the relationship between water surface temperature (WST) and standard length (SL) across the nodes of the Clupeiformes phylogenetic tree spanning 150 Myr of evolutionary history, and across the full global distribution of these fish. To evaluate the relationship between WST, SL and the species ability to move over aquatic systems we inferred the historical distance and speed of fish historical movement in a threedimensional space, using a novel phylogenetic approach (the GeoModel^30^; Methods). This model estimates the *posterior* distribution of the estimated ancestral geographical locations for all nodes in a time-calibrated phylogenetic tree – allowing us to have a measure of the distance of movement per-time unit (speed). Then, we evaluated the effect of SL and dispersal abilities on Clupeiformes speciation rates. As our approach provides information on the rate of WST change over evolutionary history, we can uniquely seek to, not only, understand how species respond to the magnitude of climate change but also how they respond to the rate at which climate has changed (how fast) over long time scales. Studying species responses to the rate of climate change is now more pertinent than ever given the alarming accelerating-rates of heating of the oceans^31^ and because species and populations respond differently when faced with a fast or slow change in their environment^32–33^.

If SL reductions under global warming decrease the ability to move and increase the probability of speciation (Fig. 1, Scenario 1), we expect to observe a negative relationship between SL and WST over both evolutionary history and across extant species (Fig. 1a); a positive relationship between dispersal abilities and SL (Fig. 1b); and a negative effect of SL on dispersal abilities and speciation rates (Fig. 1c and d). In opposition, if SL reductions under global warming increase the ability to move but decrease the probabilities of speciation (Fig. 1, Scenario 3), we expect to observe a negative relationship between dispersal abilities and SL (Fig. 1h); and a positive effect of SL and dispersal abilities on speciation rates (Fig. 1i and j). We cannot make any inference if there is no relationship between SL, dispersal abilities, and speciation rates (Fig. 1e – g, Scenario 2). Finally, if the rate of climate change can additionally modulate species responses then we should find a significant effect of the rate of WST change on both the rate of SL evolution and the speed of species movement.

## SL and WST over current and historical time

We studied the relationship between fish SL and WST over their extant geographic distribution using the phylogenetic variable rates regression model^34^ (Methods). This approach enables the simultaneous estimation of both an overall relationship between SL as a function of WST across extant species, and any significant shifts in the rate of SL evolution that apply to the phylogenetically structured residual variance in the relationship. We also included the type of migration (diadromous and non-diadromous) as an additional binary variable in the regression, as previous studies show that diadromous fish are larger on average^29^. We used a novel Bayesian approach that allows the estimation of regression coefficients while sampling the WST data within each species. With this approach we effectively evaluate the effect of WST on SL while considering the temperature variability over the entire native distributional range of each species (Methods). Results show that WST have a significant negative effect on SL across the current geographic distribution of Clupeiformes (Fig. 2a; *P*_MCMC_ = 0.001). This reveal that smaller Clupeiformes are found in warmer WST, supporting the “temperature-size rule”^35^. Diadromous species were significantly larger than non-diadromous species on average (Supplementary Table; *P*_MCMC_ = 0). Additionally, the variable rate regression did not detect any significant shifts in the rate of SL evolution, and fish SL was better explained by Brownian motion on the scaled phylogeny according to the Pagel’s Lambda (λ) parameter (Fig. 2a Supplementary Table 1).

**Figure 2.**
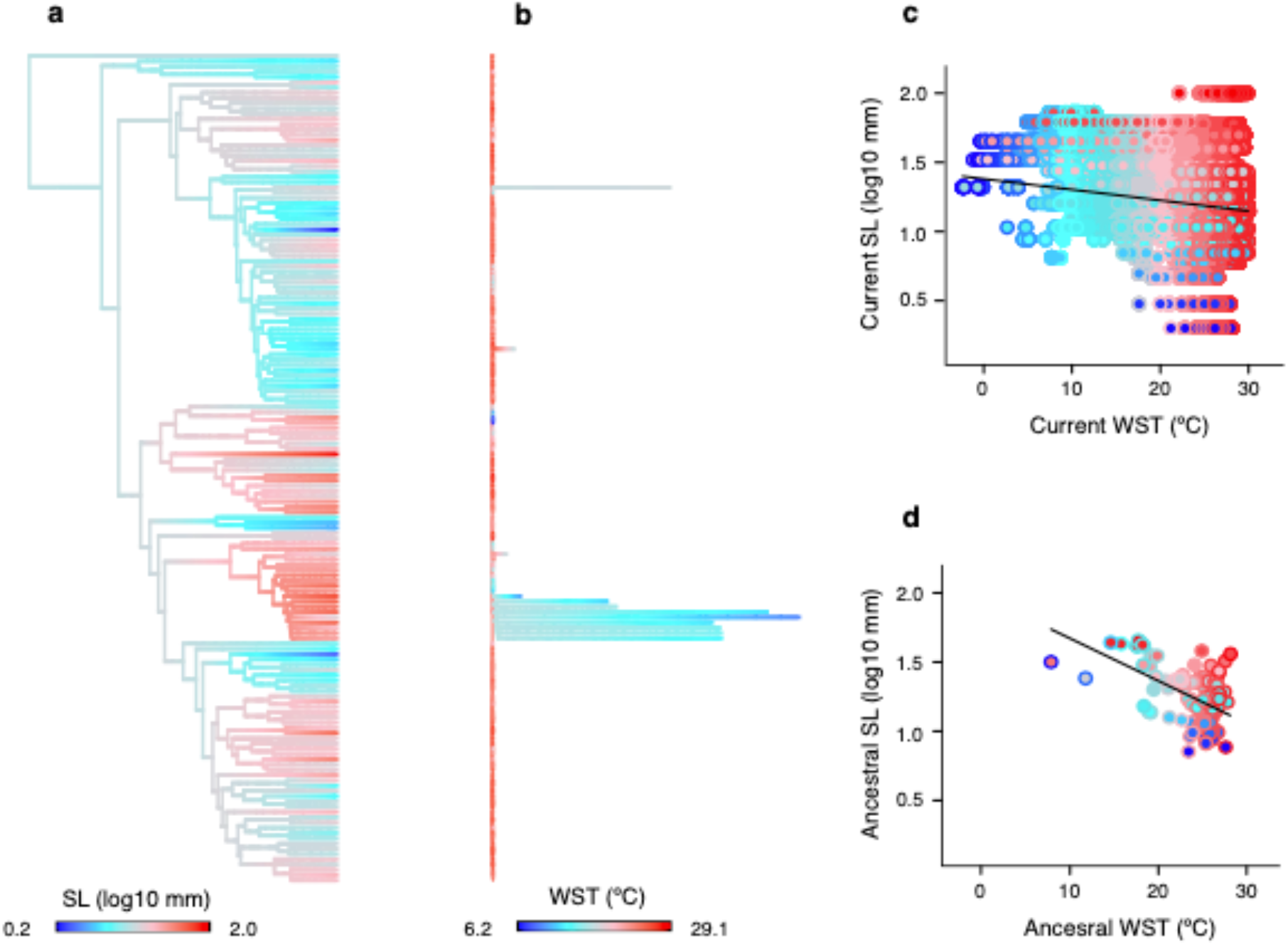
Clupeiformes evolved smaller size in warmer temperatures for million years and in recent times. **a.** Clupeiformes phylogenetic time tree with branches scaled according to (**a**) the l-model for SL, and (**b**) the variable rate regression for WST. Longer branches shows rates significantly higher than the constant-background rate (scaled in more than 95% of the *posterior* distribution). Branch colours show the ancestral states for SL and WST, estimated on the rate-scaled trees. **c.** Bayesian phylogenetic generalized least squares sustain that SL and WST are negatively correlated across extant species (*P*_MCMC_ = 0.001; *n*= 158,000). The black line represents the posterior mean slope of the phylogenetic regression, which was estimated while sampling within species WST data. **d.** Bayesian generalized least squares shows a significant negative correlation between the ancestral SL and WST values across nodes (*P*_MCMC_ = 0; *n*= 157 phylogenetic nodes; black line). Fill point gradient colours represent size values and outline point gradient colours represent WST values.

To study the relationship between fish size and temperature in the deep past, we evaluated the relationship between the SL and WST of ancestral fish and their environments, which comprises a temporal window of ~150 Myr. To conduct this analysis we, firstly, inferred the ancestral states of SL across nodes of the λ-scaled phylogeny (Fig. 2b; Methods). Secondly, we inferred the ancestral WST across nodes of the rate-scaled phylogeny (Fig. 2c) obtained from the variable rate regression between WST and absolute latitude across the 21,895 occurrence records (Methods; Supplementary Table 2). We found a significant negative association between the ancestral SL and WST (Fig. 2d; *P*_MCMC_ = 0), which support that Clupeiformes evolved smaller sizes under warmer SWT for over 150 Myr (Fig. 2d). Finally, the variable rate regression for WST indicates that the lower rate of temperature change at which Clupeiformes have survived is 0.00069 °C Myr^−1^, while the upper rate is 0.36 °C Myr^−1^ (6.9E-8 and 3.6E-6 °C per decade, respectively). These historical rates of WST change, given our data and approach, are far lower than the average rates of global warming that the planet is experiencing in the last decades; 0.07 °C per decade since 1880 to 1981, and 0.18 °C per decade since 1981 (according to the NOAA 2019 Global Climate Summary). Understanding how fish have responded to these rates of historical temperature change can provide insight of the effect that the current increasing rates of global warming will have on fish biodiversity.

## SL and dispersal abilities

The geographic analyses support a model with significant variation in the speed of fish movement across phylogenetic branches (Supplementary Table 3). This implies that the current spatial diversity of Clupeiformes have been assembled by species dispersal at variable speeds over the oceans for over 150 Myr (Fig. 3a). The average total distances taken by Clupeiformes, from the location of the most recent common ancestor (MRCA), range from 8,745 km to 55,590 km. Moreover, the speed at which these species dispersed ranges from 71 km Myr^−1^ to 536 km Myr^−1^. We evaluated the effect of SL on the total distance moved for each species from the root of the phylogenetic tree (pathwise distance; Methods), and the median of the branch-specific speed of movement along the path that links the MRCA with extant species (pathwise speed; Methods). These relationships were evaluated using Bayesian phylogenetic regression models that include a sample of 1,000 pathwise distances and speeds for each species in the estimation of regression coefficients (Methods). Results show that SL correlates positively with both the pathwise distances and the pathwise speed of movement (Fig. 3b and c; Supplementary Table 4 and 5, respectively). There were no significant differences in either the mean pathwise distances nor the pathwise speed of movement travelled by diadromous and non-diadromous species (Supplementary Table 4 and 5). These results demonstrate that smaller fish have had a reduced ability to disperse through water bodies over their evolutionary history. Therefore, smaller fish may find it hard to track suitable temperatures over geological time, thus making them more prone to extinction.

**Figure 3.**
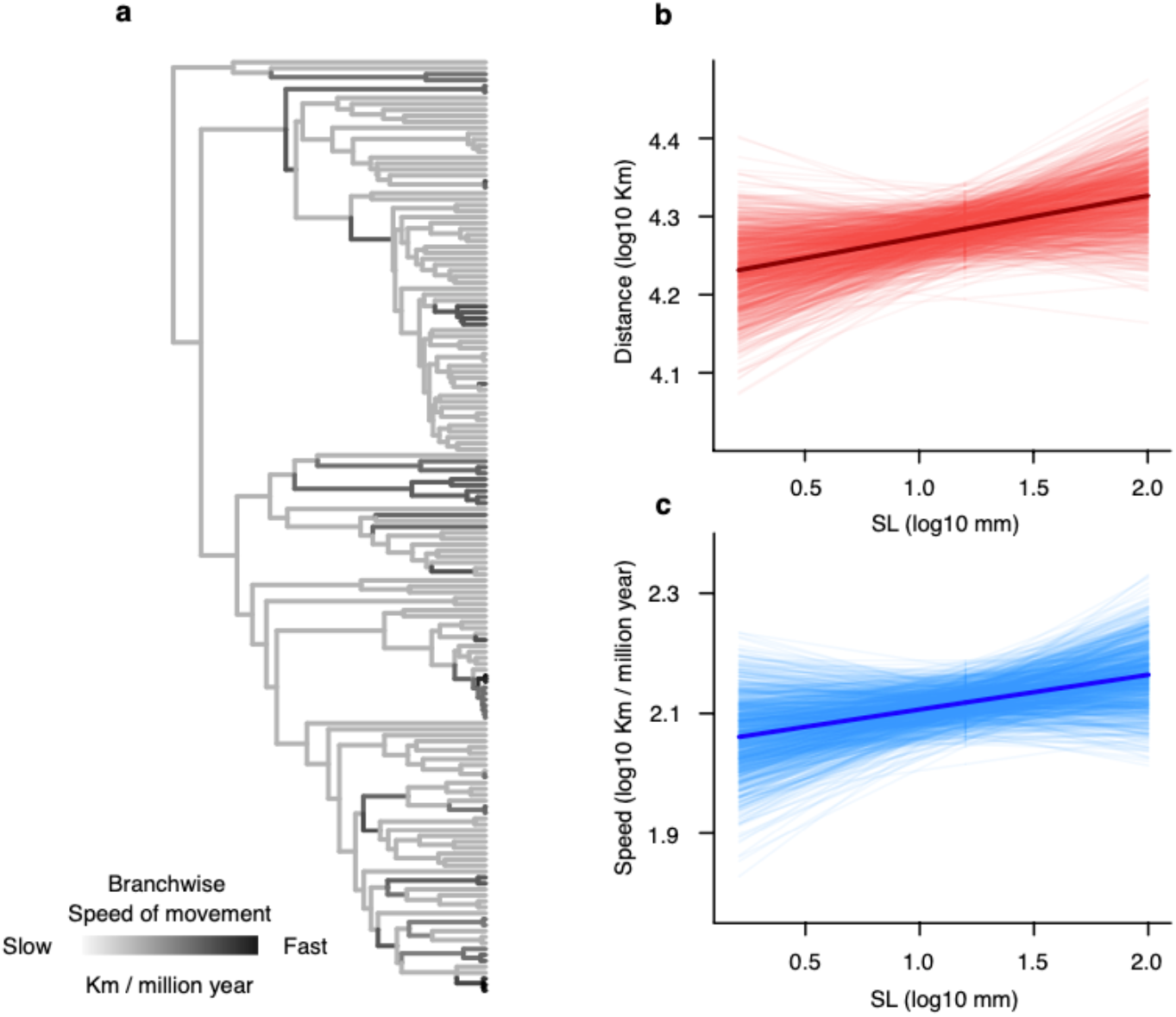
Fish dispersal abilities depend of their size. **a.** Clupeiformes phylogenetic tree with branches coloured according to the speed of movement. **b.** Bayesian phylogenetic generalized least squares show that pathwise distance correlates positively with SL (Bayes Factor > 5; *n*= 157,000). **c.** SL has also a significant positive effect on pathwise speed of movement (Bayes Factor > 10; *n*= 157,000). Lighter lines represent the posterior distribution of phylogenetic slopes and the darker lines the *posterior* mean.

## Fish response to the historical rate of WST change

We evaluated the effect that the rates of WST change may have on both the rates of SL evolution and the speed of movement across all branches of the Clupeiformes phylogeny, using Bayesian GLS regressions (Methods). All branchwise rates were estimated by dividing the scaled branches (with λ-model for SL, the variable rate regression model for WST, and the variable rate Geo Model for speed) with original branch lengths measured in time. The rate of WST change had a positive effect on both the rate of SL evolution and the speed of fish movement (*P*_MCMC_ = 0, Fig. 4a, b), meaning that the SL of Clupeiformes have evolved rapidly, and they have dispersed faster when the temperature of the oceans have changed at higher rates.

**Figure 4.**
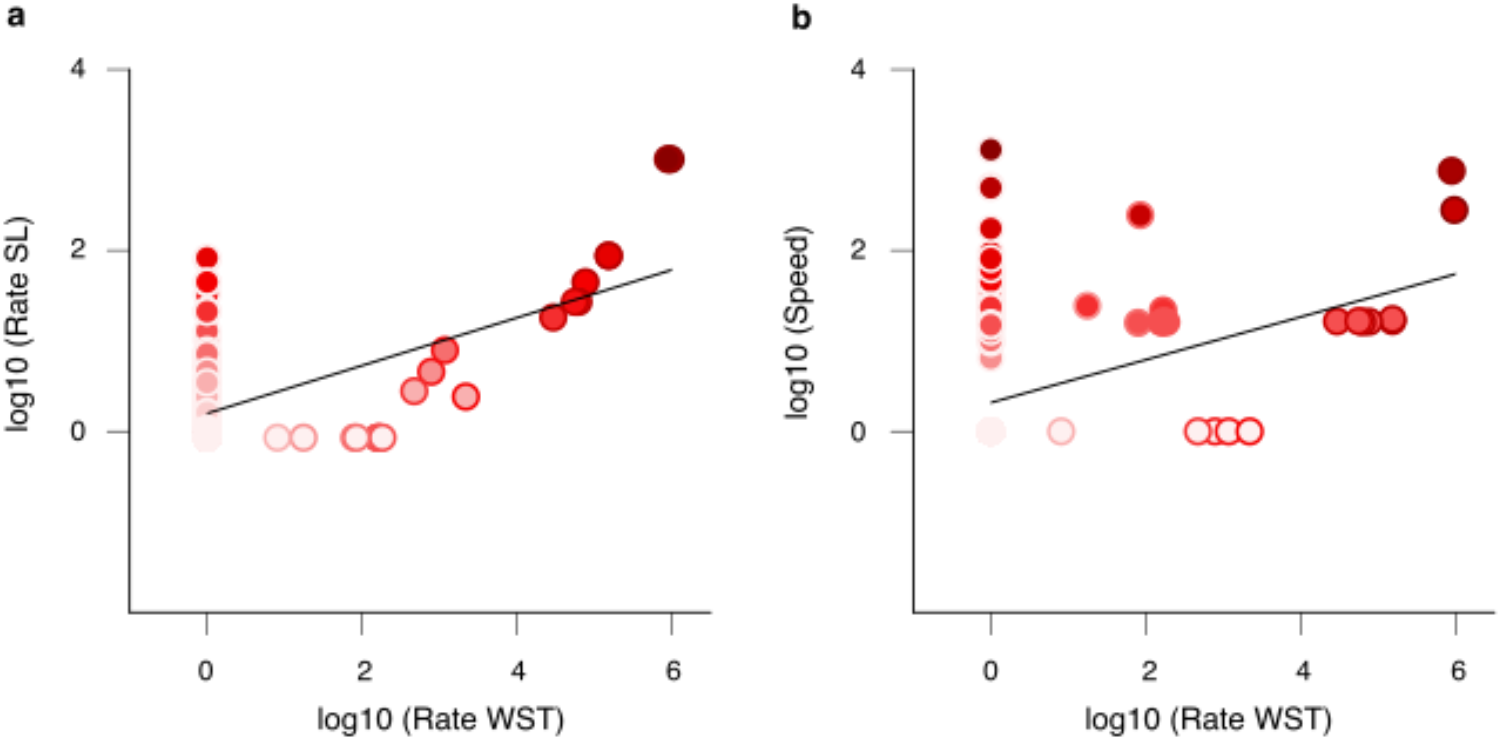
Clupeiformes have evolved rapidly and moved faster when temperature changed at higher rates. **a.** Bayesian generalized least squares support that the branchwise rates of SL evolution are positively correlated with the branchwise rates of WST change (*P*_MCMC_ = 0; *n*= 312 phylogenetic branches). **b.** The branchwise speed of fish movement are also positively correlated with the branchwise rates of WST change (*P*_MCMC_ = 0; *n*= 312 phylogenetic branches). Point fill colours represent the branchwise rates of SL evolution and the branchwise speed of species movement. Point outline colours represent the branchwise rates of WST change.

**Figure 5.**
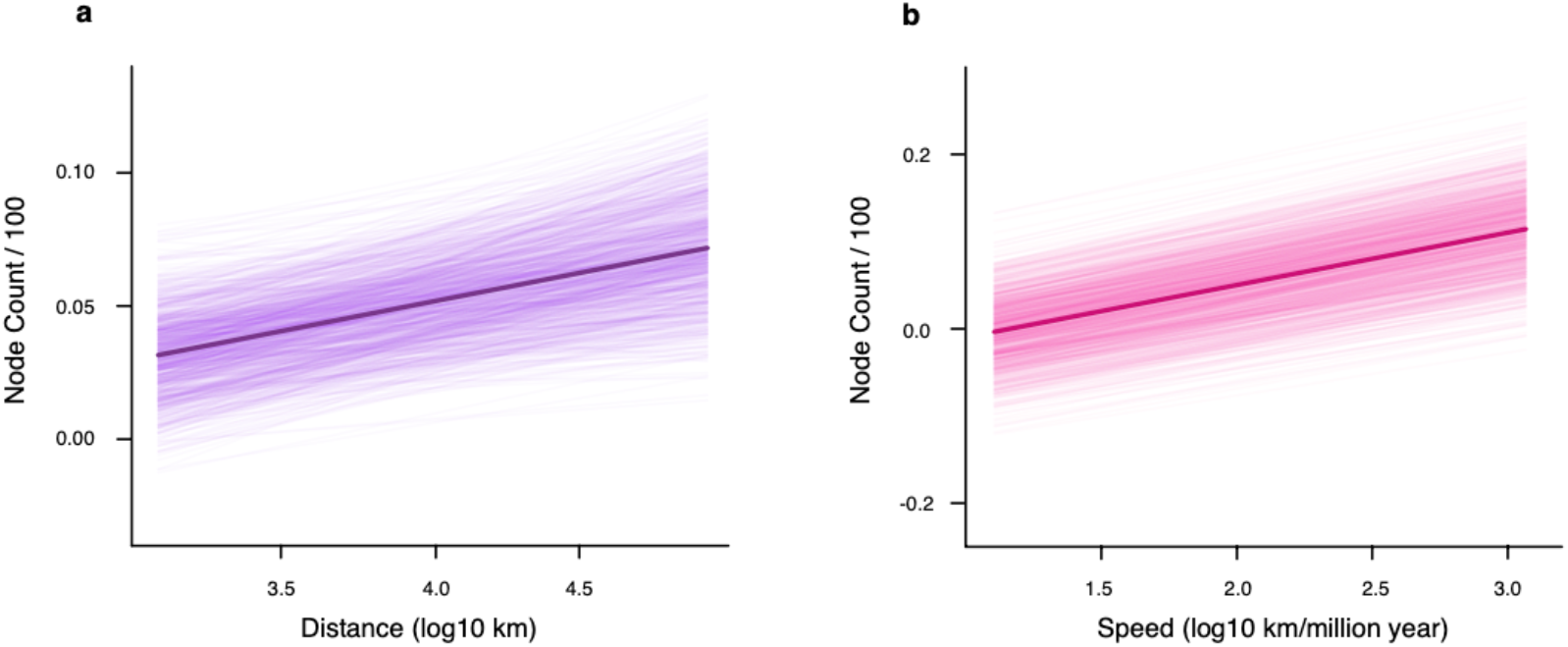
Clupeiformes with lower dispersal abilities have lower probabilities of originate new species. **a - b.** The Bayesian phylogenetic generalized least squares show that the pathwise distance of movement and the pathwise speed of movement has a positive effect on speciation (*P*_MCMC_ = 0.01 and 0, respectively; *n*= 157,000). Lighter lines show the *posterior* distribution of slopes and dark lines shows the *posterior* mean slopes. These slopes were estimated while sampling the pathwise distance and speed within species (Methods).

## Effect of SL and dispersal abilities on speciation rates

We evaluated the relationship between speciation rates and the dispersal ability and SL of Clupeiformes. We used Bayesian phylogenetic regressions models that include the uncertainty in parameter estimation and samples of dispersal abilities within species (Methods). Our results show that the independent additive effect of pathwise distance and pathwise speed were significant (*P*_MCMC_ = 0.01 and 0 respectively; Supplementary Table 6) – species that move longer distances and faster were more likely to originate new species. SL did not have a significant effect on speciation, when its independent additive effect or their interaction with dispersal ability was evaluated (Supplementary Table 6). These results suggest that fish SL, by its positive association with dispersal ability, has an indirect effect on speciation rates. The speciation rates of smaller fish that move slowly are lower than the speciation rates of their larger counterparts that moved faster and larger distances. A scenario of smaller fish under global warming may cause the loss of fish attributes that promotes the generation of biodiversity.

## Conclusion

Global change poses a double jeopardy for fish body size, as both overfishing^36^ and climate drives populations towards smaller sizes. The phenomena of fish shrinking when facing hotter waters is general in the evolutionary history of Clupeiformes and over their entire worldwide geographic distribution. Provided that smaller fish adapted to warmer conditions are less capable to disperse and in turn less able to originate new species, the scenario of global warming could limit their possibilities to find optimal environments to live and their capacity to buffer their increasing extinction risk by the process of speciation. Furthermore, Clupeiformes fish living in the present are the survivors of a long evolutionary history under variable rates of temperature change. They have responded to such historical changes by SL adaptation and dispersal but such evolutionary process have never involved the current accelerating rates of heating of the water bodies. It is probable that Clupeiformes will face an increasing risk of extinction.

## Methods

### Data

Analyses were performed on a time-calibrated phylogeny of 158 Clupeiformes species. This phylogeny was obtained from The Fish Tree of Life^37^. We used the maximum Standard Length (SL) in mm, for these 158 species, obtained from FishBase and the FAO Species Catalogue for clupeoid fishes^38^. The maximum SL was used because of three reasons. First, maximum SL is preferred over mean SL because fishes have indeterminate growth^39^. Second, it is a more stable measure of size in teleost to compare museum and collection samples. Third, and most important, individuals that are commonly larger than the population average and are outside the central distribution of size, are likely the individuals that allow the species to shift their geographic ranges^4^. 21,895 georeferenced occurrences (Supplementary Figure 1) were obtained from marine and freshwater bodies (i.e., rivers and lakes) from Aquamaps (https://www.aquamaps.org/) and the IUCN (https://www.iucnredlist.org/) respectively. We obtained the geographic locations (within the native range) of 116 species available in Aquamaps, and locations within the polygon of distribution for 42 additional species available in the IUCN. To obtain the geographic locations from the IUCN, we sampled 100 random locations within each species polygon. All georeferenced occurrences were matched with information of water surface temperature (WST). For marine species we used the mean annual sea surface temperature (SST) estimated from the Aquamaps database. For freshwater species, the mean annual air temperatures estimated from the WordClim database (https://worldclim.org/) were used as a first-order proxy of the water surface temperature of the freshwater bodies^40–42^. By maximizing the number of temperature records per species, instead of using single estimates (e.g. mean temperature, or temperate at the geographic centroid) allow us to produce more precise estimates of both the ancestral locations and the ancestral thermal environments of Clupeiformes. Finally, information about the type of migration for each species (diadromous, non-diadromous) was obtained from Bloom et al^29^.

### Inferring ancestral locations

From the geographic locations within each species in the Clupeiformes phylogeny, we inferred the ancestral geo-distribution in a continuous, threedimensional space. Ancestral locations were estimated for each phylogenetic node using the Geo Model^30^ in the computer program BayesTraits 3.0^43^. This model estimates the *posterior* distribution of ancestral locations measured in longitude and latitude, while sampling across all location-data within species, and considering the spherical nature of Earth. This natural assumption of the Earth as a spherical object avoid the erroneous calculation of distances between the inferred ancestral locations due to the non-continuity of the longitude scale. When based on a time-calibrated phylogeny, the Geo Model simultaneously estimate the speed of species movement across each branch that links pairs of phylogenetic nodes (branchwise speed of movement). The ancestral locations across phylogenetic nodes are estimated while considering the continuous variation in dispersal ability of each ancestral species - ranging from species quiescence (no movement), through constant movement in direct proportion of the passage of time, to fast species movement. Estimation of the branchwise speed of species movement are based on the variable rates model^44^ which detects shifts away from a background rate of evolution in continuous traits (expected under Brownian motion) in whole clades or individual branches. We also include data of the geographic locations of two Clupeiformes fossils, one for the crown group of Engraulidae and another for the crown group of Dorosoma. These fossils information was obtained from The Fish Tree of Life^37^.

We ran four MCMC chains for 250,000,000 iterations, sampling every 50,000 iterations, and discarding 200,000,000 as burn in. These procedures were conducted based on the Brownian motion (BM) model and the Variable Rates (VR) model (Supplementary Table 3). The final sample includes 1,000 posterior locations for each phylogenetic node. We selected the model that fit the data better by means of Bayes factors (*B*), using the marginal likelihoods estimated by stepping stone sampling. *B* is calculated as the double of the difference between the log marginal likelihood of the complex model and the simple model. By convention, *B* > 2 indicates positive evidence for the complex model, *B* = 5–10 indicates strong support and *B* > 10 is considered very strong support. We excluded the species *Denticeps clupeoides* from the Geo Model analyses because its pathwise distance and speed of movement obtained from previous analyses were outliers, which can bias the inferences made from further analyses.

### Pathwise distances and speed of species movement

In order to obtain the total distance that each species have historically dispersed through the oceans and rivers – starting from the location of the root of the Clupeiformes phylogenetic tree - we calculated the distances dispersed across each phylogenetic branch (branchwise distances) and then we summed these distances along the path that links the root with extant species (pathwise distances). The branchwise distances were calculated using the disCosine function in the geosphere R package^45^. This method calculates the great circle distance (the shortest distance) between two points on a sphere measured in kilometres using the spherical law of cosines, which works for calculating these distances at both large and small scales. We calculated the branchwise distances for every location in the posterior sample, meaning that we have 1,000 distances for every branch in the tree, and therefore, 1,000 pathwise distances for each species in the tree. With this approach we have the historical distance dispersed for each species, considering the uncertainty in ancestral locations estimates. In order to have a measure of the speed at which each species in phylogeny have dispersed over historical time, we calculated the branchwise speed of movement in km per Myr - diving the branchwise distances by the branch length of the time-calibrated tree. We also calculate the speed of movement for all the posterior sample of branchwise distances, and then we calculated the median speed of movement in the path that links the MRCA with extant species. Finally, we have 1,000 measures of the historical speed of movement for each species which include the uncertainty in ancestral location estimates.

### Phylogenetic regressions

To evaluate the expected relationships between SL, WST, pathwise distance, pathwise speed of movement, and speciation rates, we performed Phylogenetic Generalized Least Squares regression models (PGLS) with Bayesian inference which allowed us to consider the uncertainty in both, parameters estimation and within species data. We consider the uncertainty within species by using the samples of data for WST, georeferenced, pathwise distances, and speed of movement for each species. Under this approach, the MCMC samples the regression parameters and the data within species simultaneously, integrating the uncertainty of both factors in the results. All Bayesian regression were done in the computer program BayesTraits 3.0.

We, first, conducted a multiple phylogenetic regression to evaluate the relationship between SL, WST and type of migration, including the sample of WST within species. We compared the BM, Lambda model (LA), and Ornstein-Uhlenbeck model (OU) for these regressions. We also evaluated the variation in the rate of SL evolution using the variable rates (VR) regression model^34^. The VR regression model enables the simultaneous estimation of both an overall relationship between SL as a function of WST and type of migration, and any shift in rate that apply to the phylogenetically structured residual variance in the relationship. The VR regression model identify heterogeneity in the rate of evolution along phylogenetic branches (branchwise rates) by dividing the rate into two parameters: a background rate parameter (σ^2^_b_), which assumes that changes in the trait of interest are drawn from an underlying BM process, and a second parameter, *r*, which identifies a branch-specific rate shift. A full set of branchwise rates are estimated by adjusting the lengths of each branch in a time-calibrated tree (stretching or compressing a branch is equivalent to increasing or decreasing the phenotypic rate of change relative to the underlying Brownian rate of evolution). Branchwise rates are defined by a set of branch-specific scalars *r* (0 < *r* < ∞) that scale each branch to optimize the phenotypic rate of change to a BM process (σ^2^_b_ × *r*). If phenotypic change occurred at rates faster than the background rate, along a specific branch of the tree, then *r* > 1 and the branch is stretched. Rates slower than the background rate are detected by *r* < 1 and the branch is compressed. If the trait evolves at a constant rate along a branch, then the branch will not be modified (that is, *r* = 1).

Second, in order to estimate the rates of WST change through the Clupeiformes phylogeny, we conducted a Bayesian VR regression between WST and latitude (comparing it with the BM, LA, and OU regression models; Supplementary Table 2). We included the sample of WST and latitude within each species in regression analyses. Then, we obtained the consensus rate-scaled tree for WST. This consensus rate-scaled tree considers the median value of the branches scaled in more than 95% of the MCMC sample. We ran four MCMC chains for 300,000,000 iterations, sampling every 250,000 iterations, and discarding 150,000,000 as burn in. This consensus tree was also used in the estimation of the ancestral WST at phylogenetic nodes using the package phytools^46^. By using the consensus rate-scaled tree we ensure to consider the variation in the rate of SWT change when ancestral states are estimated.

Third, we evaluated the relationship between the pathwise distance with SL and type of migration, and between the pathwise speed with SL and type of migration. We included in the phylogenetic regressions the sample of species data for the pathwise distance and speed of movement, comparing regressions fitted with the BM, LA, OU, and VR model. We ran four MCMC chains for 51,000,000 iterations, sampling every 50,000 iterations, and discarding 1,000,000 as burn in. We conducted these regression using the BM, LA, OU, and VR model (Supplementary Table 4 and 5).

Fourth, we evaluated the relationship between speciation rates with pathwise speed, SL, pathwise distance, and WST - including the sample of data for pathwise distances and speed of movement. We used tip-specific estimates of speciation rates to evaluate the regression between speciation rates and the multiple explanatory variables. Among the recommended non-model-based tip-rate metrics to study the correlates of speciation rates (i.e. inverse of equal splits [ES], node density [ND] and the inverse of terminal branch length [TB])^47^ we based our interpretations on the node density along the phylogenetic paths, divided by the age of the phylogeny (100 Myr after excluding *Denticeps clupeoides*). Our choice is based on the fact that ND is the least influenced metric by potential biases and sources of uncertainty associated with branch length estimation from empirical data^48^ - ND capture the average speciation rate over the entire phylogenetic path and weight equally all branch lengths along the paths. We did not use the tip-rate speciation metric estimated from time-varying birth-death diversification models owing to the striking uncertainty in the speciation rates values when they are estimated from phylogenies with extant species only^49^, and due to the erroneous inference of the general diversification patterns when the variation in rates of sequence evolution are not properly considered in time-tree inference^50^. Additionally, we used PGLS regression models to evaluate regression-coefficients-significance because PGLS-ND has the highest statistical power when compared with PGLS-ES and PGLS-TB^47^. Furthermore, PGLS allow us to evaluate the simultaneous effect of multiple explanatory variables whose effect on speciation rates can be modelled as a linear or non-linear function. This last point is of upmost importance for our objective because there are expected interactions between the main explanatory variables (e.g. pathwise speed and SL, WST and SL) and also because there are statistical complications associated with estimating interactions without including quadratic terms (i.e. non-linear functions between the independent and explanatory variables)^51^. Our full PGLS-ND regression model is described by the following equation: ND ~ Speed + SL + Distance + WST + Speed^2^ + SL^2^ + Distance^2^ + WST^2^ + (Speed * SL) + (Distance * SL) + (WST * SL). Then, we reached the simpler reduced PGLS-ND regression model based on strict criteria: we removed the single most non-significant regression-coefficient from the full regression model, then we reiterated this procedure across every simpler regression until we get the regression with significant covariates only. We conduced these regression analyses comparing the BM and LA model. The final regression is in Supplementary Table 6. We ran 51,000,000 iterations, sampling every 50,000 iterations, and discarding the first 1,000,000 iterations as burn in. Regression coefficients were judged to be significant according to a calculated *P*_MCMC_ value for each posterior of regression coefficients. For cases in which <5% of samples in the posterior distribution crossed zero, this indicates that the coefficient is significantly different from zero.

### Non phylogenetic regressions

We applied Bayesian GLS regressions to evaluate the relationship between the branchwise rates of SL evolution, the branchwise speed of movement and the branchwise rates of temperature change. We obtained these branchwise rates and speed of movement using the rate-scaled branches as dividend and the original branch lengths (measured in time) as divisor. Specifically, we divided the branches from the LA-scaled consensus tree for SL, the VRLA-scaled consensus tree for WST, and the VR-scaled tree for geographic occurrences. Additionally, we regressed the ancestral SL on the ancestral WST inferred at each node of the Clupeiformes phylogeny. For these ancestral state reconstruction made with package phytools, we used the scaled-trees with the model that fit the data better, i.e., LA-scaled tree for SL and the VRLA-scaled tree for WST.

We conducted the Bayesian non-phylogenetic GLS regressions in BayesTraits by setting the Pagel’s Lambda parameter to zero, which discard the phylogenetic covariance of the data values. We ran 51,000,000 iterations, sampling every 50,000 iterations, and discarding the first 1,000,000 iterations as burn in. Regression coefficients were judged to be significant according to a calculated *P*_MCMC_ value for each posterior of regression coefficients. For cases in which < 5% of samples in the posterior distribution crossed zero, this indicates that the coefficient is significantly different from zero.

## Code availability

All analyses in this study were done using BayesTraits version 3 available at http://www.evolution.rdg.ac.uk/BayesTraitsV3/BayesTraitsV3.html

## Acknowledgments

We thank to Ciara O’Donovan for her help and advice with the Geo Model analyses, and to Andrew Meade for helping with the computer cluster at University of Reading. Devin D. Bloom shared data of Clupeiformes. JA-LL was supported by ANID FONDECYT postdoctoral grant N° 3200654 and ANID FONDECYT regular grant N° 1201506. CBCA was supported by ANID FONDECYT initiation grant N° 11180897 and he was partially supported by Nucleo Milenio INVASAL funded by the Chilean government program Iniciativa Científica Milenio. CEH and RJR was supported by ANID FONDECYT regular grant N° 1170815 and 1201506. MMR was supported by ANID FONDECYT regular grant N° 1200843. CV was supported by a Leverhulme Trust Research Project Grant (RPG-2017-071) and a Leverhulme Trust Research Leadership Award (RL-2019-012).

## Author contributions

JA-LL and CBCA formulated and developed the overarching idea and research goals. JA-LL and CV designed the methodology and created the statistical models. JA-LL implemented the computer codes and applied the statistical analyses of data. CEH provided computational support for data analysis. OIM and RJR contributed to obtain the dataset and making of figures. JA-LL wrote the original draft and figures. CBCA, CV and MMR critically reviewed the original draft. All authors made comments, suggestion and editions to the last draft.

## Competing interests

The authors declare no competing interests.

## Correspondence and request for materials

should be addressed to JA-LL.

## Supplementary Figures

**Supplementary figure 1.**
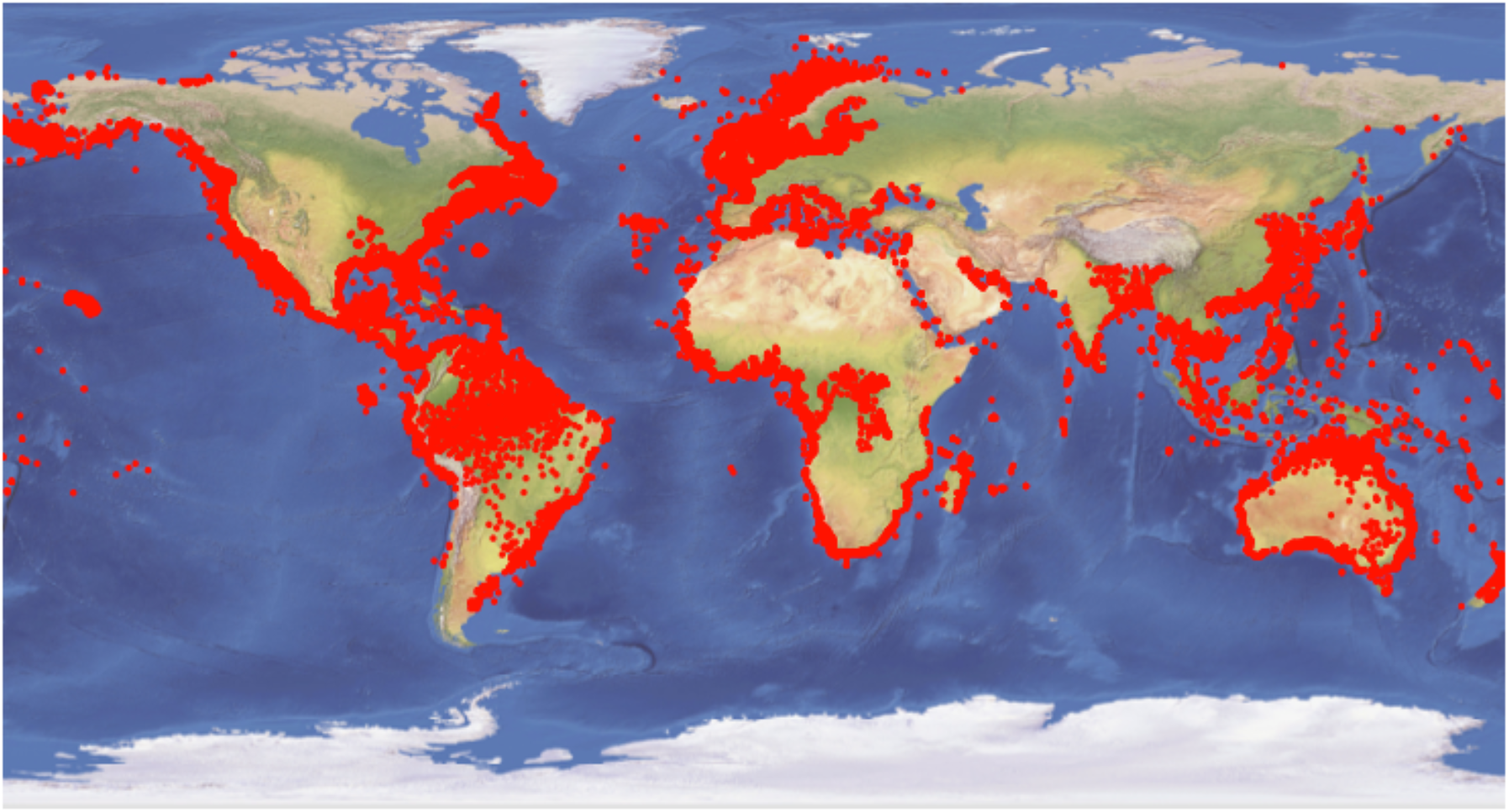
Geographic distribution of Clupeiformes species used in this study. Red dots represents the geographic occurrences obtained from Aquamaps and the IUCN which comprises 21,895 datapoints for 158 species. This dataset was used for the ancestral locations inference and to obtain data of environmental temperature.

## Supplementary Tables

**Table 1.** Evolutionary model fitting for the regression that evaluate the effect of type of migration and water surface temperature (WST) on fish standard length (SL). Data analysed includes the maximum SL and samples of WST, within the native range, for each species. The log Marginal Likelihood (Marginal Lh), estimated by stepping stone sampling, provides the models support given the data and priors. More positive values support a given model, where differences >1 indicates positive evidence; differences between 2,5 - 5 indicates strong support; and differences > 5 indicates very strong support for a model over the other. BM = Brownian Motion, LA = Lambda, OU = Ornstein-Uhlenbeck, VR = Variable Rate, VRLA = Variable Rate and Lambda.

| SL Phylogenetic Regression Model | Marginal Lh.<br>BM | Marginal Lh.<br>LA | Marginal Lh.<br>OU | Marginal Lh.<br>VR | Marginal Lh.<br>VRLA |
| --- | --- | --- | --- | --- | --- |
| $SL \sim \alpha + \beta_1(\text{Diadromous}) + \beta_2(\text{WST})$ | -59.11 | **8.09** | -19.84 | -16.29 | 8.13 |

**Table 2.**
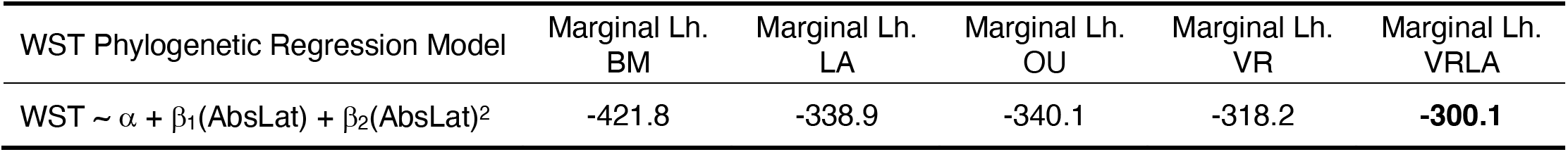
Evolutionary model fitting for the regression that evaluates the effect of absolute latitude on WST. Data analysed includes a sample of WST and absolute latitude (AbsLat) within the native range of each species. The log Marginal Likelihood (Marginal Lh), estimated by stepping stone sampling, provides the models support given the data and priors. More positive values support a given model, where differences >1 indicates positive evidence; differences between 2,5 - 5 indicates strong support; and differences > 5 indicates very strong support for a model over the other. BM = Brownian Motion, LA = Lambda, OU = Ornstein-Uhlenbeck, VR = Variable Rate, VRLA = Variable Rate and Lambda.

**Table 3.**
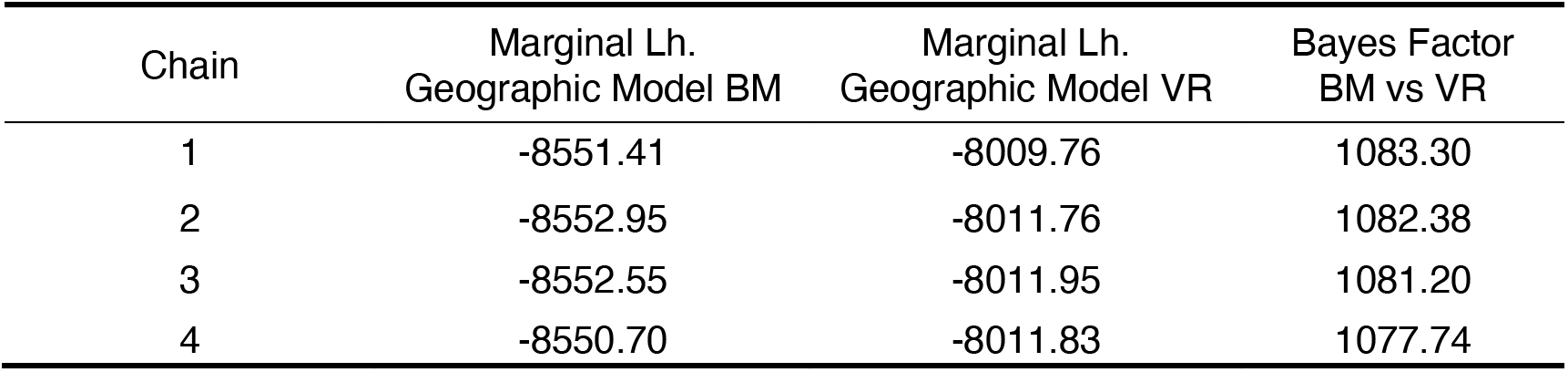
Geographic model (Geo Model) fitting for Clupeiformes georeferenced data. The Geo Model estimate the longitudes and latitudes across the nodes of the phylogenetic tree by means of Bayesian inference. These coordinates are estimated onto a three dimensional cartesian coordinates system which were modelled using Brownian motion (BM) – the rate of location change across the tree is constant. We also allowed the rate of location-change to vary across phylogenetic branches by fitting the Variable Rate model (VR). The log Marginal Likelihood (Marginal Lh), estimated by stepping stone sampling, provides the models support given the data and priors. More positive values support a given model, where differences >1 indicates positive evidence (Bayes Factor > 2); differences between 2,5 - 5 indicates strong support (Bayes Factor 5 – 10); and differences > 5 indicates very strong support for a model over the other (Bayes Factor > 10).

| Chain | Marginal Lh.<br>Geographic Model BM | Marginal Lh.<br>Geographic Model VR | Bayes Factor<br>BM vs VR |
| --- | --- | --- | --- |
| 1 | -8551.41 | -8009.76 | 1083.30 |
| 2 | -8552.95 | -8011.76 | 1082.38 |
| 3 | -8552.55 | -8011.95 | 1081.20 |
| 4 | -8550.70 | -8011.83 | 1077.74 |

**Table 4.** Evolutionary model fitting for the regression that evaluate the effect of SL and type of migration on the speed of fish movement. The log Marginal Likelihood (Marginal Lh), estimated by stepping stone sampling, provides the models support given the data and priors. More positive values support a given model, where differences >1 indicates positive evidence (Bayes Factor > 2); differences between 2,5 - 5 indicates strong support (Bayes Factor 5 – 10); and differences > 5 indicates very strong support for a model over the other (Bayes Factor > 10). BM = Brownian Motion, LA = Lambda, OU = Ornstein-Uhlenbeck, VR = Variable Rate, VRLA = Variable Rate and Lambda.

|  | Marginal Lh.<br>BM | Marginal Lh.<br>LA | Marginal Lh.<br>OU | Marginal Lh.<br>VR | Marginal Lh.<br>VRLA |
| --- | --- | --- | --- | --- | --- |
| Distance $\sim \alpha + \beta_1(\text{SL})$ | 138.71 | 133.99 | 134,99 | **148.04** | 140.46 |
| Distance $\sim \alpha + \beta_1(\text{SL}) + \beta_2(\text{Diadromous})$ | 133.03 | 125.89 | 129.43 | 138.4 | 136.29 |
| Distance | 123.57 | 118.07 | 117.37 | 128.77 | 126.12 |

**Table 5.** Evolutionary model fitting for the regression that evaluate the effect of SL and type of migration on the distance of fish movement. The log Marginal Likelihood (Marginal Lh), estimated by stepping stone sampling, provides the models support given the data and priors. More positive values support a given model, where differences >1 indicates positive evidence (Bayes Factor > 2); differences between 2,5 - 5 indicates strong support (Bayes Factor 5 – 10); and differences > 5 indicates very strong support for a model over the other (Bayes Factor > 10). BM = Brownian Motion, LA = Lambda, OU = Ornstein-Uhlenbeck, VR = Variable Rate, VRLA = Variable Rate and Lambda.

|  | Marginal Lh.<br>BM | Marginal Lh.<br>LA | Marginal Lh.<br>OU | Marginal Lh.<br>VR | Marginal Lh.<br>VRLA |
| --- | --- | --- | --- | --- | --- |
| Speed $\sim \alpha + \beta_1(\text{SL})$ | 102.67 | 96.99 | 99.71 | **135.16** | 109.77 |
| Speed $\sim \alpha + \beta_1(\text{SL}) + \beta_2(\text{Diadromous})$ | 93.07 | 91.97 | 89.61 | 113.02 | 101.47 |
| Speed | 83.13 | 82.79 | 81.37 | 94.13 | 87.81 |

**Table 6.** Phylogenetic regression model for Node Density (ND) obtained after reducing the full model ND ~ Speed + SL + Distance + WST + Speed^2^ + SL^2^ + Distance^2^ + WST^2^ + (Speed * SL) + (Distance * SL) + (WST * SL). The log Marginal Likelihood (Marginal Lh), estimated by stepping stone sampling, provides the models support given the data and priors. More positive values support a given model, where differences >1 indicates positive evidence (Bayes Factor > 2); differences between 2,5 - 5 indicates strong support (Bayes Factor 5 – 10); and differences > 5 indicates very strong support for a model over the other (Bayes Factor > 10).

|  | Marginal Lh.<br>BM | Marginal Lh.<br>LA |
| --- | --- | --- |
| $\text{ND} \sim \alpha + \beta_1(\text{Speed}) + \beta_2(\text{Distance})$ | **524.02** | 517.17 |

## Notes

### Competing Interest Statement

The authors have declared no competing interest.

